# African swine fever virus – variants on the rise

**DOI:** 10.1101/2022.09.07.506908

**Authors:** Jan H. Forth, Sten Calvelage, Melina Fischer, Jan Hellert, Julia Sehl-Ewert, Hanna Roszyk, Paul Deutschmann, Adam Reichold, Martin Lange, Hans-Hermann Thulke, Carola Sauter-Louis, Dirk Höper, Svitlana Mandyhra, Maryna Sapachova, Martin Beer, Sandra Blome

## Abstract

African swine fever virus (ASFV), a large and complex DNA-virus circulating between soft ticks and indigenous suids in sub-Saharan Africa, has made its way into swine populations from Europe to Asia. This virus, causing a severe haemorrhagic disease (African swine fever) with very high lethality rates in wild boar and domestic pigs, has demonstrated a remarkably high genetic stability for over 10 years. Consequently, analyses into virus evolution and molecular epidemiology often struggled to provide the genetic basis to trace outbreaks while few resources have been dedicated to genomic surveillance on whole-genome level. During its recent incursion into Germany in 2020, ASFV has unexpectedly diverged into five clearly distinguishable linages with at least ten different variants characterized by high-impact mutations never identified before. Noticeably, all new variants share a frameshift mutation in the 3’ end of the DNA polymerase PolX gene O174L, suggesting a causative role as possible mutator gene. Although epidemiological modelling supported the influence of increased mutation rates, it remains unknown how fast virus evolution might progress under these circumstances. Moreover, a tailored Sanger sequencing approach allowed us, for the first time, to trace variants with genomic epidemiology to regional clusters. In conclusion, our findings suggest that this new factor has the potential to dramatically influence the course of the ASFV pandemic with unknown outcome. Therefore, our work highlights the importance of genomic surveillance of ASFV on whole-genome level, the need for high-quality sequences and calls for a closer monitoring of future phenotypic changes of ASFV.

## Introduction

It is widely accepted today that most virus populations consist of a variety of genetic variants rather than one clonal virus. The emergence of these virus variants is driven by the virus specific mutation rate [1], which depends on multiple factors including mode of replication, fidelity of polymerases, the availability of repair mechanisms, as well as selection. Together, these two factors are responsible for the speed with which evolution progresses (evolutionary rate), demonstrated by the emergency of new virus variants [2]. While some viruses evolve very fast and new variants develop quickly, impressively demonstrated during the recent SARS-CoV2 pandemic [3], other viruses demonstrate a high degree of genetic stability and evolve very slowly. One example for the latter is the African swine fever virus (ASFV) [4,5].

This large and complex DNA virus has been first described in Kenya in 1921 [6], where it is transmitted in an ancient sylvatic cycle between warthogs and soft ticks of the genus Ornithodoros [7,8]. Since 2007 the virus was translocated to Eurasia and spread in wild boar and domestic pig population. While the African warthogs remain largely asymptomatic after infection [9], the virus is highly lethal to domestic pigs [10,11] and Eurasian wild boar [11–13]. Although distantly related viruses have been identified in amoebae and some degree of similarity to irido- and poxviruses has been shown, no closely related viruses are known today [5]. Therefore, ASFV was only recently grouped into the phylum Nucleocytoviricota and, because it is the only known member of its family Asfarviridae and the genus Asfivirus [5], is still considered a mystery in modern virology.

The ASFV genome, a single molecule of covalently closed double stranded DNA with a size of up to 190 kbp [14], has a remarkably high genetic stability. Modern virus strands show a very high degree of nucleotide sequence identity to viral elements integrated in the soft tick genome dated to at least 1.46 million years [15]. This observation is supported by recent analyses of the ASFV strain introduced into Georgia in 2007. Despite over ten years of epidemic circulation, the virus strain has accumulated only very few overall mutations and even less affecting viral genes [16,17].

When ASFV was introduced into the wild boar population of eastern Germany in 2020 [18], whole genome sequencing revealed an ASFV strain similar to the strains known to circulate in western Poland including a mutation within the O174L gene, coding for ASFV DNA repair polymerase X [19,20]. This insertion of a tandem repeat was utilized together with other mutations in K145R, MGF 505-5R and the intergenic region between 173R and I329L as genetic marker to trace outbreak clusters in affected Polish counties. [21] While this discovery was no doubt interesting, no evidence for differences in the virus phenotype were observed at that point. What came as a surprise was the subsequent detection of numerous ASFV variants in Germany characterised by high impact mutations affecting known ASFV open reading frames (ORFs) that have never been described before. While some of the changes affect regions of the viral genome that could be linked to potential immune modulators or virulence factors, the influence of most mutations remains unknown. Therefore, we wanted (i) to investigate in more detail what might underlie this new genetic variability, (ii) to utilize this newly emerged genetic variance for molecular epidemiology, and (iii) to identify consequences of our findings on the epidemiological level using models.

The present manuscript summarizes the canon of all these investigations and suggests that the previously described mutation in the O174L gene coding for ASFV polymerase X has led to an increased mutation rate and thus higher evolutionary rate culminating in the emergence of the viral variants.

## Material and Methods

### DNA extraction

#### For routine diagnostics

Field samples were extracted using the QIAamp® Viral RNA Mini kit (Qiagen, Hilden, Germany) or the NucleoMagVet kit (Macherey-Nagel, Düren, Germany) on a KingFisher® extraction platform (Thermo-Fisher-Scientific, Waltham, USA) according to the manufacturer’s instructions.

#### For next-generation sequencing

DNA was extracted from the original sample matrices using the NucleoMagVet kit (Macherey-Nagel) according to the manufacturer’s instructions. The extracted DNA was stored at −20°C until analysis. Prior to NGS analysis, dsDNA from the samples was quantified using a Nanodrop spectrophotometer (ThermoFisher Scientific).

#### For sanger sequencing

DNA was extracted from field samples using the QIAamp® Viral RNA Mini kit (Qiagen) according to the manufacturer’s instructions.

### PCR

#### Routine ASFV diagnostics

Field samples were analysed by an OIE listed ASFV specific qPCR [22] including a heterologous internal control [23] and by the commercial virotype ASFV 2.0 kit (Indical Biosciences). The latter included both a heterologous and an endogenous internal control and was carried out according to the manufacturer’s instructions. All analyses were done on a Bio-Rad C1000TM thermal cycler (BIO-RAD, Hercules, USA), with the CFX96TM Real-Time System of the same manufacturer.

### Sequencing

#### Sample selection

To identify samples suitable for shotgun sequencing, e.g. samples with a favourable ratio between host and viral genome, ASFV positive samples showing a difference of at least 5 Cq values between the ASFV target and the house-keeping gene beta actin as host genome representative (used as internal control) were chosen from the pool of routine diagnostic samples and DNA samples received from the Ukraine stored at −20° at the FLI.

#### Next-generation sequencing

For NGS, DNA was extracted from the original sample matrices using the NucleoMagVet kit (Macherey-Nagel) according to the manufacturer’s instructions (Supplementary Table S1). The extracted DNA was stored at − 20°C until analysis. Prior to NGS analysis, dsDNA from the samples was quantified using a Nanodrop spectrophotometer (ThermoFisher Scientific).

Subsequently, a minimum of 100 ng of DNA was sent to and sequenced by Eurofins Genomics. This service included preparation of a 450 bp DNA sequencing library using a modified version of the NEBNext Ultra™ II FS DNA Library Prep Kit for Illumina and sequencing on an Illumina NovaSeq 6000 with S4 flowcell, XP workflow and in PE150 mode (Illumina, San Diego, USA).

#### iSeq 100 sequencing

For Illumina iSeq 100 sequencing, DNA sequencing libraries were produced using the GeneRead DNA Library I Core Kit (Qiagen) and Netflex Dual-index DNA Barcodes (Perkin Elmer, Waltham, USA) according to the manufacturer’s instructions. Prior to sequencing, libraries were analysed on a Bionalayzer2100 (Agilent, Santa Clara, USA) using the High Sensitivity DNA Analysis kit (Agilent) and quantified using the KAPA Library Quantification Kit for Illumina® Platforms (Roche, Basel, Switzerland). iSeq 100 sequencing was performed according to the manufacturer’s instructions in 150 bp paired-end mode using an iSeq 100 i1 Reagent v2 (300-cycle) kit (Illumina).

#### MiSeq sequencing

Sample preparation was performed as described above for the iSeq100. Final libraries were sequenced on the Illumina MiSeq using the Reagent Kit v2 or v3 (Illumina) according to the manufacturer’s instructions.

#### Sanger sequencing

DNA was extracted from field samples using the QIAamp® Viral RNA Mini kit (Qiagen) according to the manufacturer’s instructions. Marker identification and genetic typing of 834 positive tested field samples was realized by amplicon generation and sanger sequencing of nine target ASFV genome regions. To this end, conventional PCR was performed using designated primers (Supplementary Table 2) and the Phusion Green Hot Start II High Fidelity PCR Master Mix (Thermo-Fisher-Scientific) according to the manufacturer’s instructions in a 25 μl reaction on a C1000 Thermo Cycler (Biorad, Hercules, USA). Subsequently, PCR reactions were sent to and analysed by Microsynth Seqlab GmbH (Göttingen, Germany) or Eurofins Genomics (Ebersberg, Germany). The service included PCR clean-up and sanger sequencing.

### Data analysis

#### Next-generation sequencing

NGS data from German field samples was analysed by mapping all reads against the ASFV Germany 2020/1 (LR899193) [18] genome sequence as reference using Newbler 3.0 (Roche) with default parameters including adapter and quality trimming. Subsequently, mapped reads were extracted and assembled using SPAdes 3.13 [24] in the mode of error correction prior to assembly with default parameters and automatically chosen K-mer length. Assembled contigs were assessed in Geneious Prime® 2021.0.1 and manually modified where necessary (especially in G/C homopolymer regions). For validation, all reads were mapped to the assembled contig using Newbler 3.0 and the sequence was corrected manually when necessary. For detection of novel ASFV variants, ASFV whole-genome sequences were aligned to the ASFV Germany 2020/1 (LR899193) [18] genome sequence as reference using MAFFT v7.450 [25] in Geneious Prime. The obtained 22 whole-genome sequences were submitted to the European Nucleotide Archive (ENA) under the project accession PRJEB55796.

#### Comparison of GTII whole-genome sequences from public databases

Sequences were downloaded from INSDC databases. To reduce the rate of calling false positive mutations due to sequencing errors, sequences were chosen due to the availability of quality parameters such as a mean coverage per nucleotide of at least 40 and aligned using MAFFT v7.450 [25] in Geneious (Supplementary Table 3). Furthermore, due to the inaccuracy of modern sequencing platforms to correctly call the number of G/C nucleotides in homopolymer stretches and frequent sequence artefacts due to low coverage at the genome ends, the extensive G/C homopolymer-regions at the 5’-end as well as the ITR regions (genome position <1000 and >190000) of the ASFV genome were excluded from the analysis.

#### Sanger sequencing

The received data from sanger-sequencing was analysed in Geneious Prime by alignment to the ASFV Germany 2020/1 (LR899193) [18] genome sequence as reference using MAFFT v7.450 [25].

#### Epidemiological modelling

We investigate data from model simulations using the software SwiFCo-rs (for technical documentation see https://ecoepi.eu/ASFWB/). The model links individual animal behaviour to the spatio-temporal structure of wild boar population over thousands of square kilometres. Hence, individual level knowledge about infection, transmission and virus genome drives the observable outcome on the landscape or population level. The model was verified, validated, and applied with different problems of ASFV epidemiology [26]. The model is developed in the Rust language and used as Python library. The latter is available from the authors upon reasonable request.

The model compiles (i) an ecological component detailing processes and mechanisms related to the ecology, sociology and behaviour of wild boar in natural free-roaming populations of the species Sus scrofa; (ii) an epidemiological ASF component reflecting individual disease course characteristics and transmission pathways including direct contact on different social scales and environmental transmission caused by ground contamination or contacts to carcasses of succumbed infected host animals; and (iii) a pseudo-genetics component manipulating inheritance of code patterns with every successful infection between two wild boar individuals. The model is stochastic in relation to all three components and parametrised using reported distributions from literature including variability and uncertainty [27].

The basic principle of transmission relates to the number of adjacent/in contact animals and carcasses using event probabilities, i.e. each infectious object provides a chance of transmission to every susceptible animal sufficiently close. The wild boar-ASF-system comprises three modes of potential transmission, i.e. between live animals of the same social group (within group transmission), between live animals of different groups (between group transmission) and between carcasses of animals succumbed to the infection and live animals (carcass-mediated transmission). Parametrisation of the modes of transmission integrates multiple sources [28–30].

The model runs on habitat maps reversely calibrated to generate spring population density according to European density models [31] and covering about 200km to the West and East of the German Polish border. Dynamic visualisations of model runs are available from https://ecoepi.eu/ASFWB/VAR. All model runs were performed on the same geographical landscape. The infection was released in the north-eastern part of the simulation landscape. Simulated spread generated westwards and southwards waves with continuous approach towards the Polish-German border.

Variant dynamics were determined by the parameter mutation probability. Whenever a transmission event occurred, the newly infected animal either inherits the variant of the source of the infection or is assigned a completely new variant not yet attached to any other individual. The variants are modelled as opaque identifiers without a genetic code. This avoids having to describe how and where a variant changed the genetic information.

The output measure per simulation was the spatial distribution of variants, and the number of variants that covered more than 100 km^2^ by varying the rate at which new variants stochastically occur. Furthermore, we estimated the probability distribution to detect exactly one out of three samples and at least 10 variants from 50 samples selected from the infectious carcasses on the German side in the first year since arrival of the simulated epidemic at the border.

## Results

### Whole genome sequencing reveals ten distinct ASFV variants in Germany

Genome analysis revealed five lineages (I-V) with a total of ten variants (Figure 1). Whole-genome sequencing was successful for 22 ASFV positive field samples representing different areas of disease introduction. They comprised of either EDTA-blood, blood-swabs or bone marrow. Twenty-two whole-genome sequences (WGS) were successfully assembled with mean coverages per nucleotide varying from 21.4 – 943.4 (Supplementary Table S1). These German ASFV sequences show a very high overall nucleotide sequence identity to other available ASFV GTII WGS from International Nucleotide Sequence Database Collaboration databases (INSDC) and clearly belong to P72 GTII (Supplementary Figure S1 A). However, through alignment to the first generated ASFV WGS from Germany (LR899193.1) [18], five lineages (I-V) with a total of ten variants were identified based on single nucleotide variations (SNV) as well as insertions or deletions (indels) of one or two nucleotides (Table 1, Figure 1 and Supplementary Figure S1).

**Figure 1.**
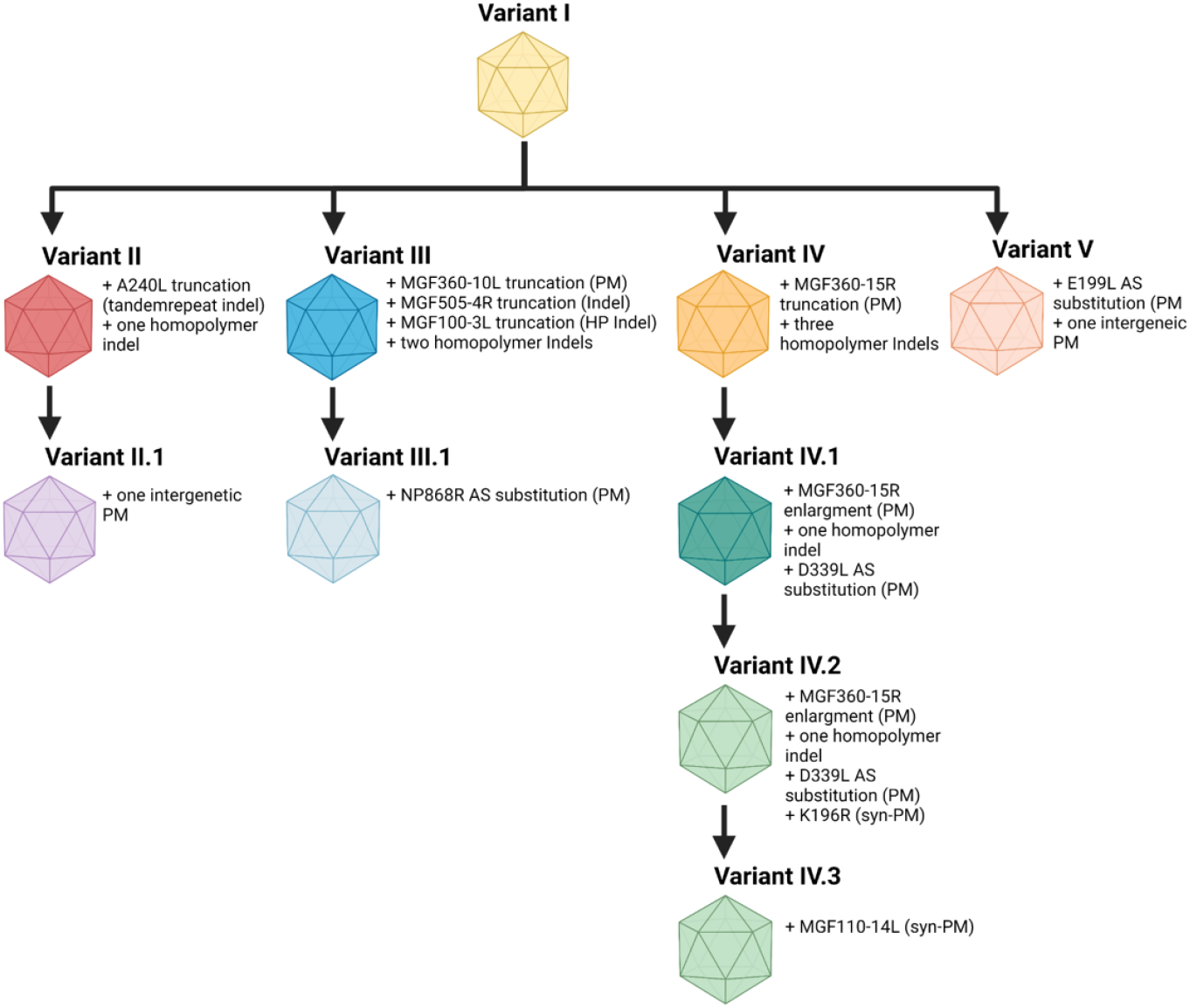
ASFV variants in Germany. Variants are characterised by insertions/deletions (indels) in homopolymer and non-homopolymer regions as well as substitutions (subst.) leading to the discrimination of five lineages and ten variants.

**Table 1.**
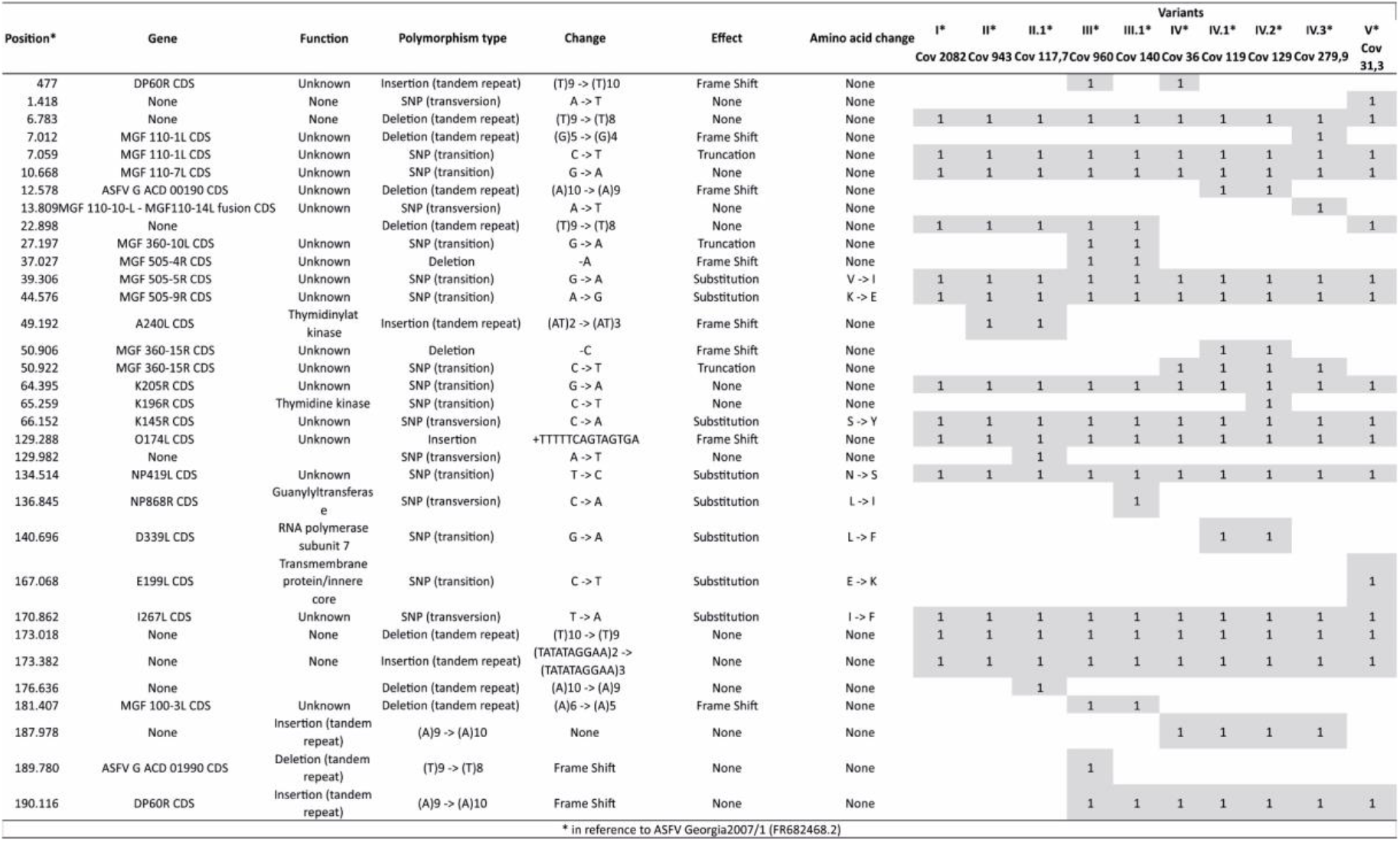
Genetic differences in German ASFV variants compared to ASFV Georgia2007/1 (FR682468.2).

### ASFV variants in Germany are characterized by 13 novel mutations affecting annotated open reading frames (ORFs)

When compared to the first generated German ASFV WGS LR899193.1 [18], the 22 WGS of German ASFV presented here are characterized by 17 novel mutation sites of which 13 affect annotated ORFs. These mutations affect five multigene family (MGF) genes, MGF110-14L, MGF360-10L, MGF505-4R, MGF360-15R, and MGF100-3L as well as DP60R, ASFV G ACD 00190 CDS, ASFV G ACD 01990 CDS, A240L, K196R, NP868R, D339L, and E199L (Table 1).

Of these 13 ORF-affecting mutations, six indels leading to frameshifts resulting in six truncations (MGF505-4R, A240L, MGF360-15R, MGF100-3L, ASFV G ACD 00190 CDS and ASFV G ACD 01990 CDS) and two nonsense mutations (stop codon integration) leading to truncation (MGF360-10L and MGF360-15R) are classified as high-impact mutations (HI-Mutation). Furthermore, three substitutions leading to one amino acid change (NP868R, D339L and E199L) and two synonymous mutations (K196R and MGF110-14L) (Figure 1 and Table 1) are classified as low-impact mutations (LI-Mutation).

### Stochastic emergence of geographic clusters of variants

We ran epidemiological model simulations starting with few infected wild boars at the location where the first cases were confirmed in Western Poland in 2019, 200km distant to previous virus circulation. Figure 2 illustrates the geographic emergence of spatial clusters of variants. The randomly emerging variants (different colours in Figure 2) formed spatially separated clusters on the German side of the border. Figure 2 a-c illustrates the temporal development of variant clusters in a single simulation run. The further the spread of the infection branches geographically the more individual variant clusters emerge. In Figure 2 d-f we show the variant map at the end of three different simulation runs using identical model parameters.

**Figure 2.**
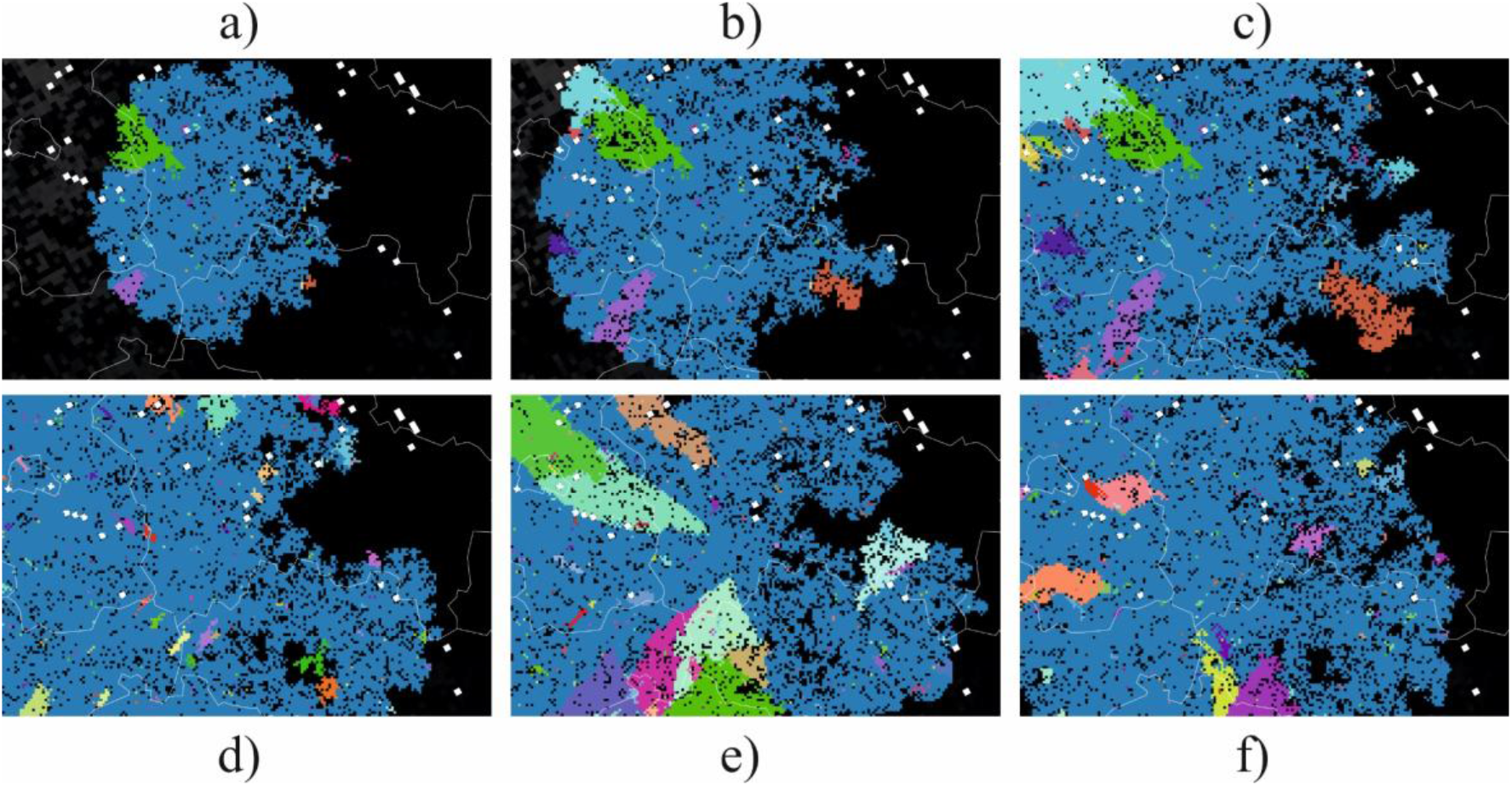
Model output as spatial snapshots with different variants differently coloured, either showing the dynamic development of infection distribution (a-c) or maping the stochastic variability of the final distribution (d-f). Pixels represent social groups of individual wild boar and lines are administrative borders.

More systematic, using model output of 100 runs, Figure 3 shows the counts of variants that formed a minimum cluster size of 100 km^2^ dependent on the parameter *mutation probability*, describing the rate of variant emergence per new animal infection (animal passage). The cluster size of 100 km^2^ was used to reflect the cluster dimensions found in Germany while excluding containment measures. The rate of variant emergence per animal passage that resulted in at least 10 variants with cluster size of 100 km^2^ was at 1.15% (Figure 3). The spatial clustering of variants in the model output does suggest such relationship between the variants found in the field.

**Figure 3.**
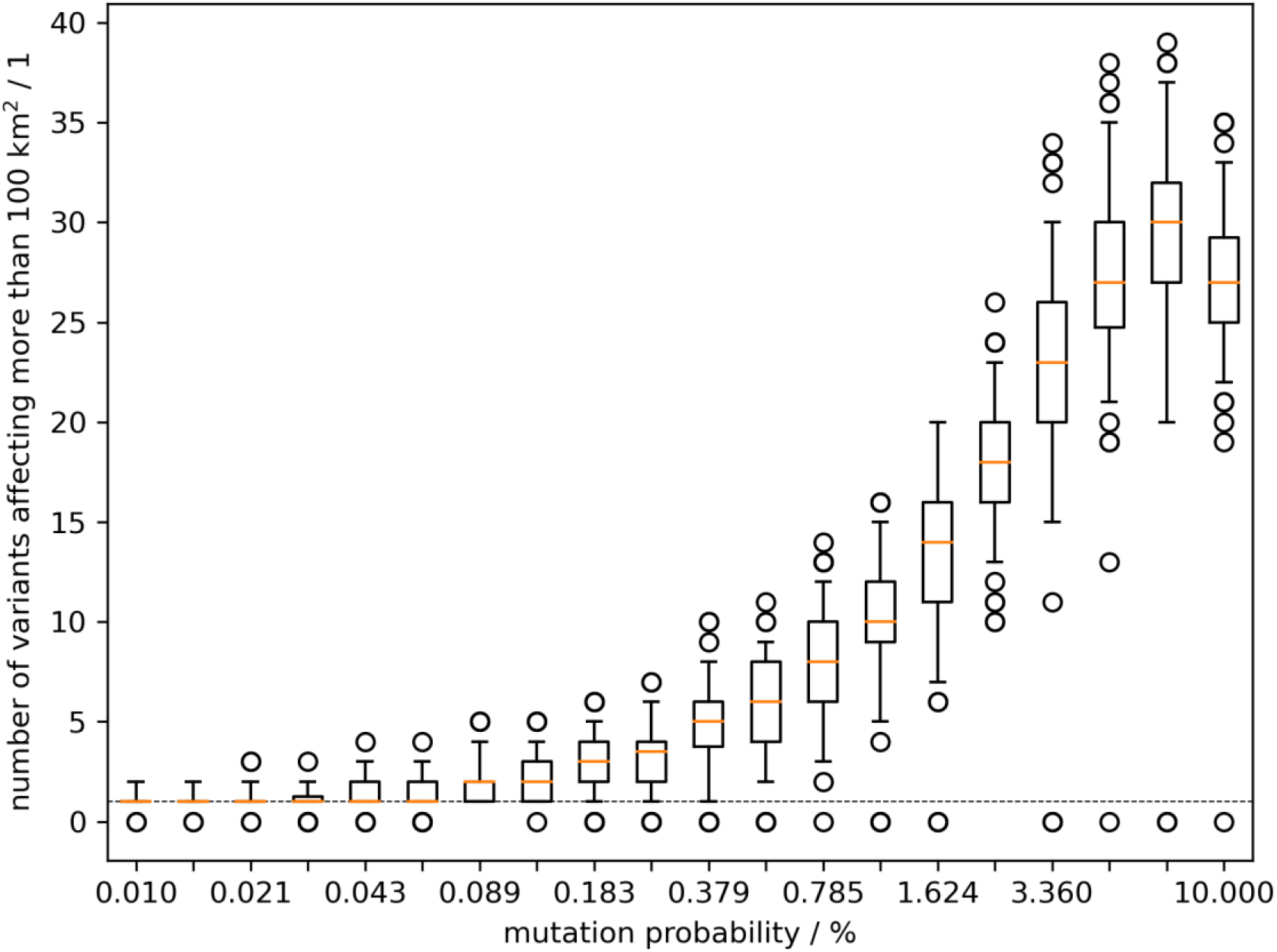
Model outcome of the number of emerging variants which affected at least 100 km^2^ of wild boar habitat. Box whisker plots summarise 100 model runs per value of the mutation parameter.

### Mutation sites can be used as markers for genomic epidemiology of ASFV in Germany

The 22 newly generated WGS as well as the previously published ASFV Germany 2020/1 (LR899193) genome sequence were used as template for PCR primer design to amplify nine different mutation-regions as genetic markers (Supplementary Table 2). In total 834 field samples were successfully assigned to one of the ten variants (Figure 1). When geographically displayed, a clear spatial clustering was detected. Variants of Lineages I and II, i.e. I, II and II.1 were found in the East Brandenburg districts of Oder-Spree (see Figure 4, LOS) and its neighboring districts Spree-Neiße (SPN, Variant I - northern part) and Dahme-Spreewald (LDS, Variant II), whereas viruses of Lineage III were detected in the more northern districts Märkisch-Oderland (MOL), Barnim (BAR) and Uckermark (UM). Variant III.1 was detected in MOL, Frankfurt (Oder) (FF) and LOS. Variants of the Lineage IV were found in the southern areas like SPN Variant IV – southern part) and, so far, represent the sole variants detected in the federal state of Saxony (Görlitz - GR, Variant IV, IV.1, IV.2, IV.3). The distribution of Variant V spans closely to the Polish border from FF to LOS. Notably, all involved German districts, with the exception of Dahme-Spreewald (LDS), share a common border with Poland.

**Figure 4.**
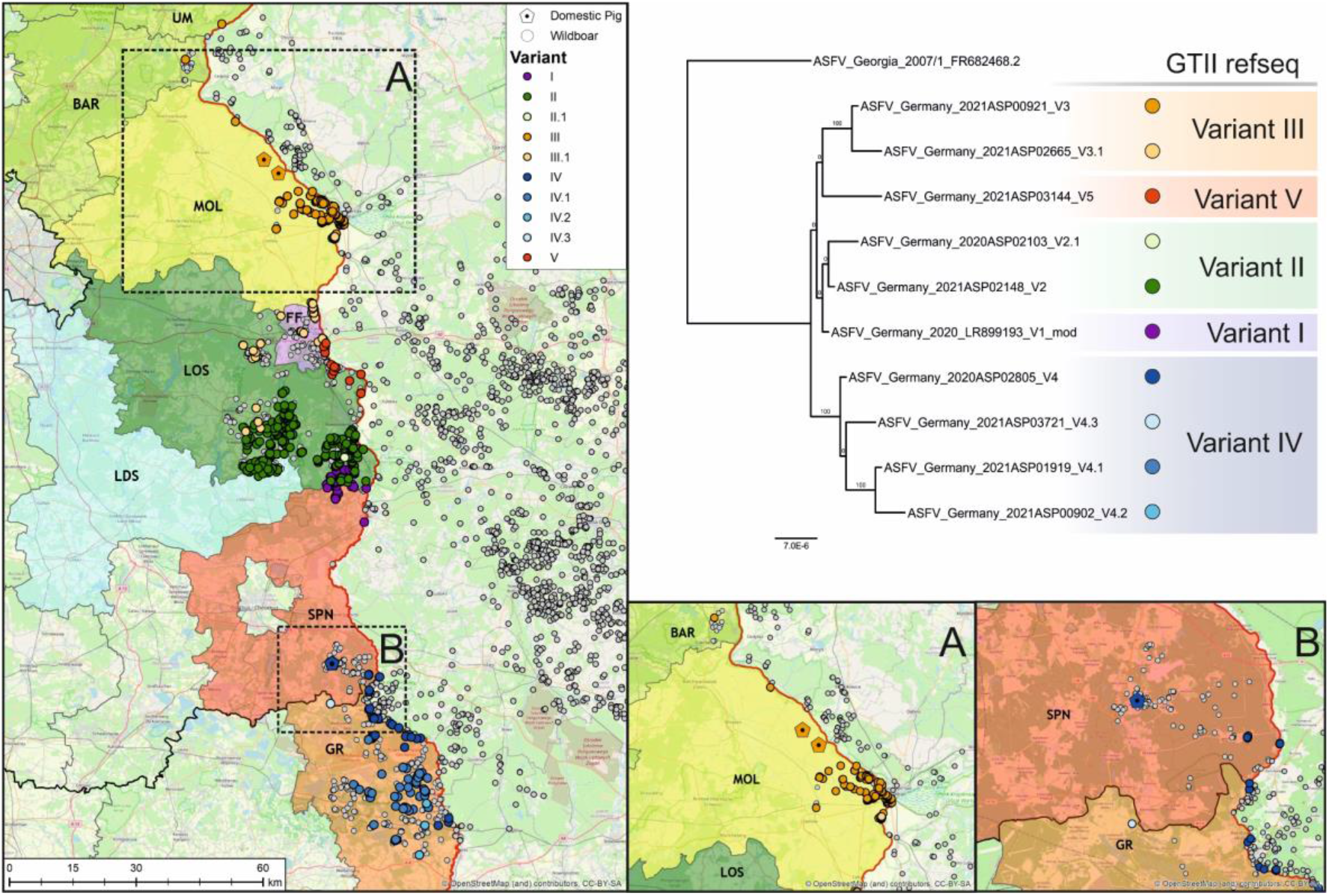
Distribution of viral variants detected in the federal states of Saxony and Brandenburg along the Polish border (left). Confirmed ASFV cases in wild boars from September 10th 2020 until August 12th 2021 are depicted as circles (white), whereas outbreaks in domestic pigs are shown as pentagons (n = 3, areas A and B). In order to facilitate the visualization of spatial ASFV clusters, variants confirmed by Sanger sequencing (n = 834) were coloured according to their assignment to one of the five lineages: Variants of Lineage I are depicted in violet, variants of Lineage II in green, variants of Lineage III in orange, variants of Lineage IV in blue, and Lineage V in red. The phylogenetic tree is provided to aid Lineage and variant understanding and was constructed using IQ-TREE version 1.6.5 [42,43] on MAFFT v7.450 [25] aligned ASFV whole-genome sequences. Standard model selection was used, resulting in the best-fit model GTR+F+R4. Statistical support of 10,000 ultrafast bootstraps using the ultrafast bootstrap approximation (UFBoot) (percentage values) is indicated at the nodes.

### Variants analysis suggest local spill-over from wild to domestic hosts

Three outbreaks in domestic pigs occurred within the study period (Figure 4 A, B). Using whole-genome sequencing, all three outbreak strains could be assigned to variants circulating in the immediate vicinity of the outbreak farms. In detail, lineage III was found in two domestic pig outbreaks in the district MOL while lineage IV was found in SPN. In all three cases, the variant was first detected in wild boar, hence an introduction from the local wild boar population is likely.

### Compared to worldwide ASFV GTII whole-genome sequences ASFV Germany shows excessive high-impact-mutations

We evaluated if the findings in Germany indicate novel and different situation regarding the frequency of high impact mutations. Altogether, 35 international ASFV WGS were compared (Supplementary Table 3). We chose 21 publicly available ASFV WGS originating from eastern and western Europe (including the first German sequence), Russia and Asia from 2007-2020 due to (I) availability and geographic distribution and (II) available sequence quality parameters e.g. mean coverage per nucleotide >50) (Supplementary Table 3). We added five WGS generated from samples of domestic pigs collected in the Ukraine from 2017-2018. Finally, we included nine WGS from Germany that represented the other viral variants found in Germany to this date. The 35 sequences were examined for their genetic variance relative to the sequence of ASFV introduced into Georgia in 2007 [32]. Due to frequent issues in sequencing the inverted terminal repeat regions and resulting variations in sequence length, only genome positions (in regard to ASFV Georgia 2007/1) from 1423-187970 were included.

In total, 131 variations were detected including 34 indels and 97 nucleotide substitutions (Supplementary Table 3). From 96 mutations affecting annotated ORFs, 81 are Low-Impact-mutations (LI-mutations) leading to amino acid substitution and 15 are High-Impact-mutations (HI-mutation) leading to stop-codon integration or frameshift. From these 15 HI-mutations, nine can be detected in German ASFV sequences (from which eight are exclusive and one is shared with other GTII sequences) (Figure 5). Thus, of all HI-mutations recorded in 35ASFV GTII WGS over 14 years 53 % (8/15) specifically occur in ten German ASFV sequences collected in about one year.

**Figure 5.**
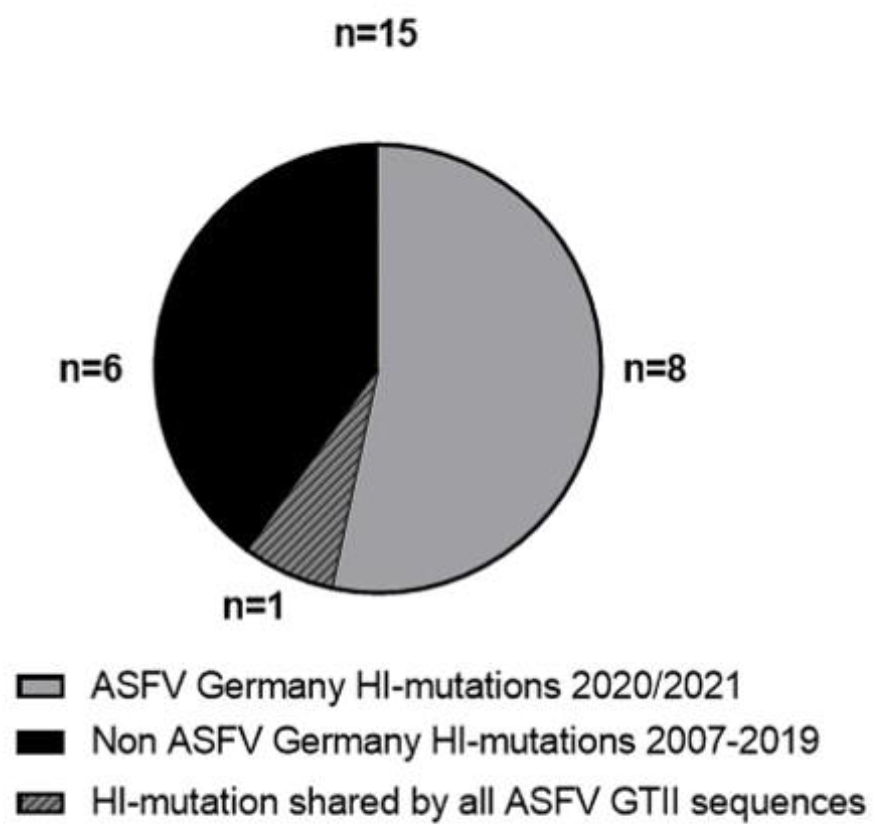
High-impact-mutations in the ASFV whole-genome sequences. Number of HI-mutations in the 23 ASFV WGS from Germany vs HI-mutations in 5 WGS from Ukraine and 20 publicly available WGS. All compared to ASFV Georgia 2007/1 (FR682468.2).

### Analysis of ASFV WGS sequences from the Ukraine in a comparable spatiotemporal scenario shows genetic variability but only few high-impact-mutations

To compare the situation in Germany to another country with similar geographical and temporal distribution of ASFV outbreaks, we analysed samples collected in 2017/2018 from domestic pig outbreaks in northern Ukraine by whole-genome sequencing. In total, WGS from five samples were successfully assembled with mean coverages per nucleotide ranging from 68.5-209.8. When aligned to the ASFV-Georgia2007/1 (FR682468.2) [32] as reference, 32 mutations were detected from which ten are LI-mutations and two are HI-mutations leading to the truncation of annotated genes (MGF300-4L, and ASFV G ACD 00270 CDS) (Supplementary Table 1 and 4). When compared with the German ASFV variants, the total number of mutations is comparable (31 for Germany and 32 for the Ukraine) but the number of novel HI-mutations is higher (eight for Germany and two for the Ukraine) (Supplementary Table 1 and 4). The different ratio of HI:LI mutations (8:23 vs. 2:30) contradicts the assumption that both ratios reflect similar mutation dynamics (Fisher exact 0.043, p < 0.045).

### Plausibility of difference in number of variants in separated virus populations

We tested on a model set up (1 variant out of 3 analysed samples. vs. 10 variants out of 50 analysed samples) whether the number of variants detected in sequencing data from the Baltics (low sample number example) was compatible with the number found in Germany (large sample number example) under the assumption that the mutation rate did not change between these settings. Figure 6 combines the model predictions for one variant detected from 3 genetically determined samples, called P(v=1|s=3), with those of 10 variants out of 50 samples, called P(v≥10|s=50). The former captures the available data of the Baltics where no variants were detected and 3 WGS of the virus were assembled (blue distribution), the latter captures the situation in Germany where 10 variants were detected using 50 genome sequences of the virus sampled during the first year after entry (orange distribution).

**Figure 6.**
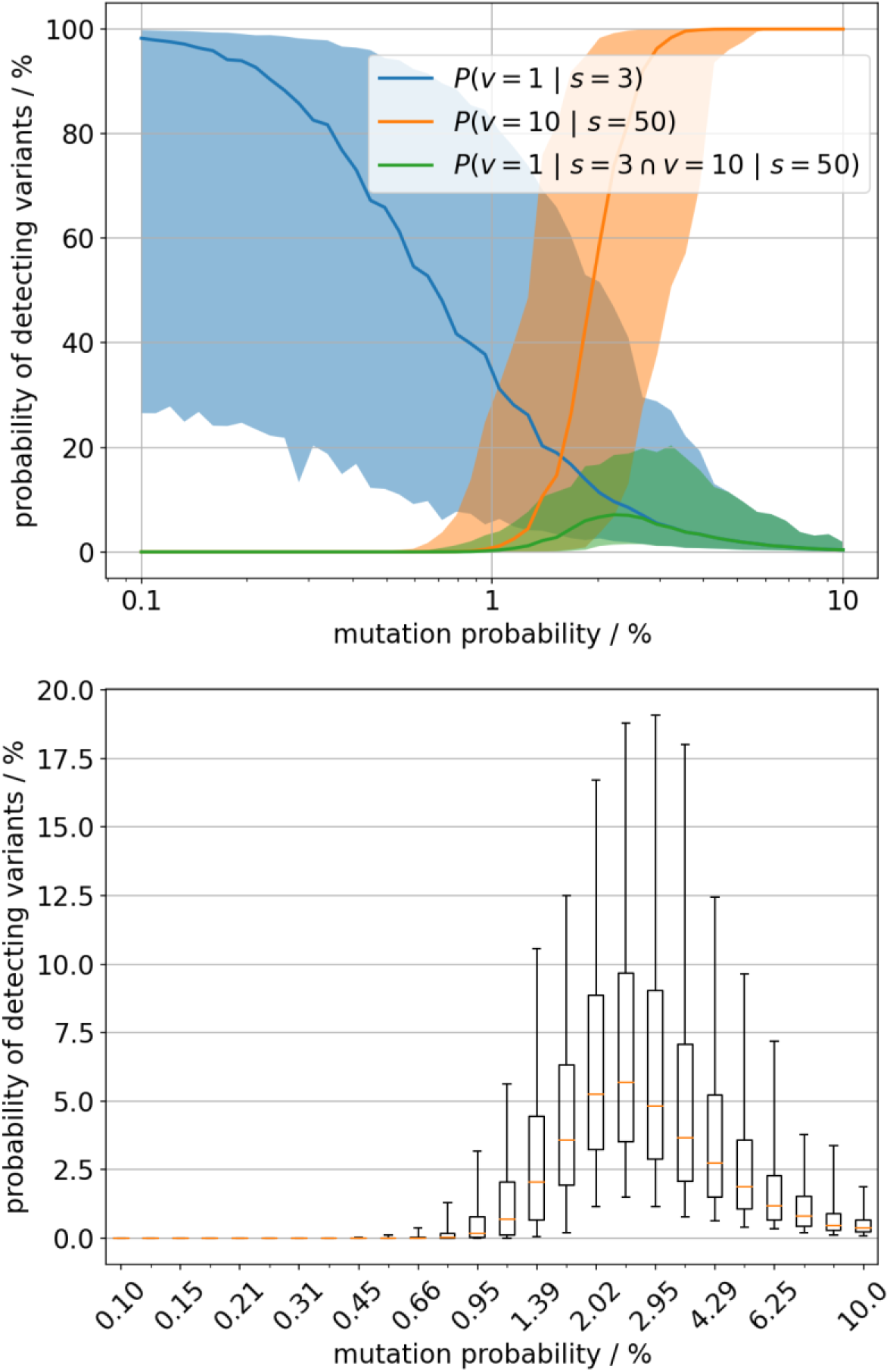
A) Likelihood of observing 1 variant out of 3 sequenced samples (blue) and 10 out of 50 sequenced samples (orange) shown by the median value (bold line) and the 90% credibility interval (shaded area). The probabilities are estimated for varying rate of variants’ emergence (x-axis, log scaled). The green graph represents the joint distribution i.e. the probability to observe both sample outcomes with constant variants’ emergence rate. B) Distributional details of the green graph i.e the joint probability.

The two sample outcomes (1/3 & 10/50) give an estimate of the situation in the past and thus we are interested in the probability of their joint occurrence in the model setting. The probability of both sequence sampling outcomes together was factually zero for large ranges of variant emergence rate (Figure 6A). The eligible range of positive variants’ emergence rate is very narrow around 2%. However, even there the probability of joint observations of both sequencing data is only about 5% in median (Figure 6B). Therefore, the model data suggested that the two sequencing scenarios more likely result from virus populations with different variant emergence rates. The simulation cannot identify the reason for the difference in variants’ emergence rate, this was addressed by structure analysis of mutation in the O174L gene of the field samples.

### A mutation in the ASFV polymerase X (O174L gene) might act as mutator and contribute to the increased number of ASFV variants

In order to identify possible reasons for the increased occurrence of ASFV variants in Germany, we surveyed the German ASFV WGS for mutations that might act as mutators, i.e. such mutations, that increases the viral mutation rate. Through alignment to other ASFV GTII sequences (FR682468.2), we detected a previously described HI-mutation shared by all German and three Polish ASFV WGS. A 14 bp duplication at pos. 129288 (in reference to ASFV Georgia2007/1 FR682468.2[32]) leads to a frameshift and truncation of the O174L gene (Figure 7A, Table 1 and Supplementary Table 3) [18–21]. This gene encodes the DNA polymerase X (PolX), a well-characterised enzyme involved in base-excision repair [33–35]. The frameshift results in a truncation by seven amino acid residues from the C-terminal end (R168-L174) as well as substitution of an additional eight residues, four of which lie within the last α-helix of the enzyme, called αF (Figure 7B). Although the conformation of this αF helix (residues 156-163) is likely preserved in the mutant owing to the conservative nature of its four amino acid substitutions, the terminal peptide connected to this helix (residues 164-167) likely adopts a different conformation in the mutant. This assumption is based on the substitution of the helix-breaking glycine-164 residue of the wild type for a helix-stabilizing leucine in the mutant (Figure 7B-C).

**Figure 7.**
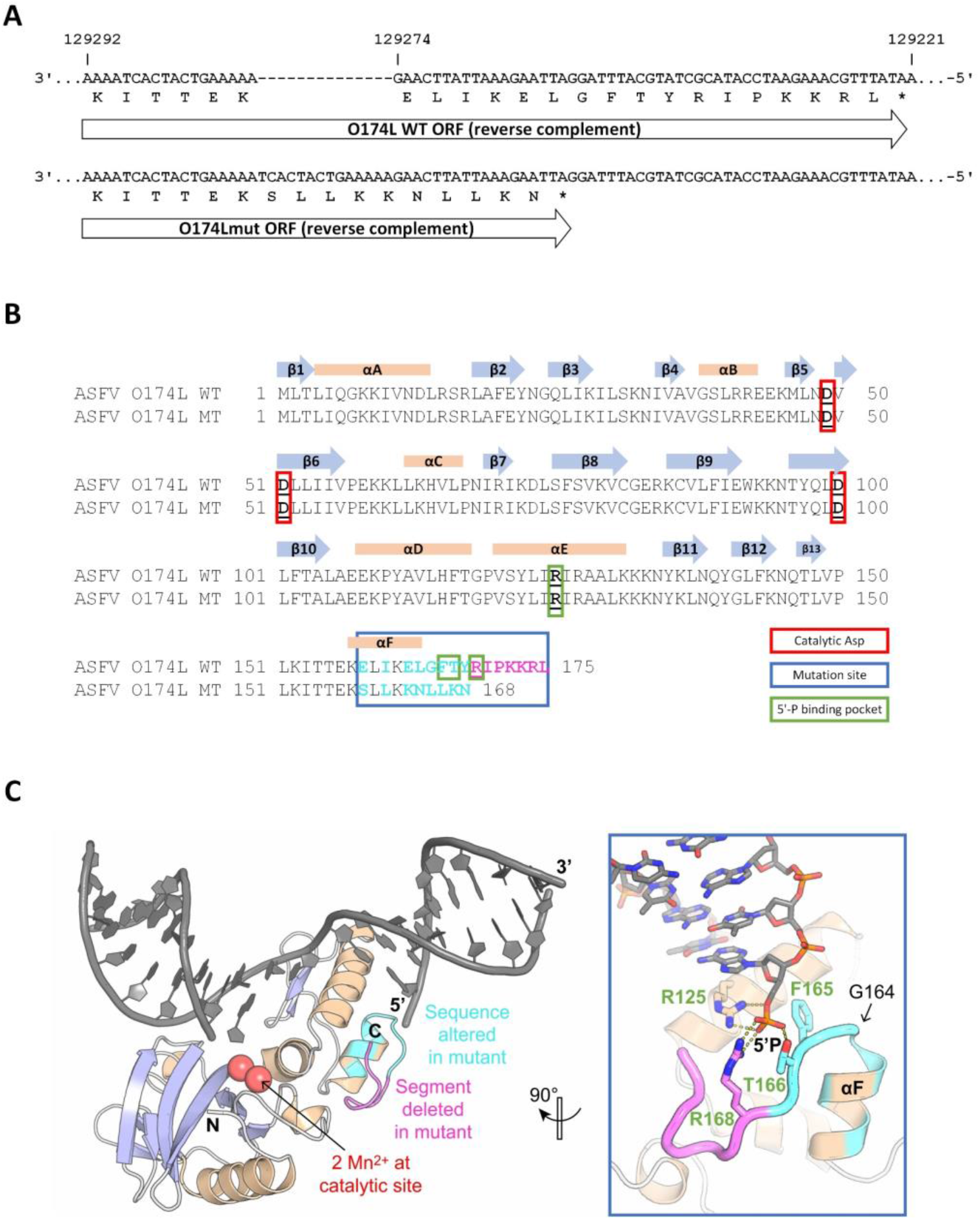
Comparison of O174L wildtype and mutant nucleotide and protein sequence and the effects of observed mutations on the wild-type ASFV PolX protein structure. Alignment of ASFV O174L wildtype and mutant nucleotide sequence (A) and protein sequence (B) including structural information from the literature [34]. Catalytic sites (red box), mutation site (blue box), amino acids forming the 5’-binding pocket (green box) and variated amino acids (red letters) are highlighted. The nucleotide alignment was done using MAFFT v7.4506 and the protein alignment using Clustal W in Geneious. (C) X-ray structure of wild-type ASFV PolX in complex with nicked DNA (PDB accession: 5HRI) (reference: PMID: 28245220). Positions with altered sequence in the mutant are coloured in cyan and positions that are missing in the mutant are coloured in magenta. The illustration was prepared with PyMol (Schrödinger, Inc.).

In the wild-type enzyme, the C-terminal region including the αF helix forms part of a positively charged pocket composed of residues R125, T166, and R168 that bind the negatively charged 5’-phosphate end of DNA substrates at single-strand breaks that are introduced into the repair sites by the viral apurinic/apyrimidinic (AP) endonuclease [34]. Whereas R125 remains unaffected by the mutation, the other positively charged residue of the pocket, R168, is lost in the deletion. It is however possible that the substitutions E162K and T166K, which introduce two new positive charges, compensate for this loss (Figure 7A-B). Therefore, the mutant PolX enzyme may still be overall functional, whereas its kinetic and thermodynamic parameters, or its substrate specificity, are likely affected.

## Discussion

Despite the extremely high genetic stability of the ASFV genome, the existence of genetic variation is not surprising and has been documented in previous studies [16,21,36]. However, the data we present in this study on ASFV variants in Germany are unexpected and show an extraordinary development that has not been described before. Within one year of ASFV spread in German wild boar resulting from multiple introductions several geographical clusters have been formed that are associated with virus sub-populations characterised by high impact mutations. The herein presented data give evidence for at least five lineages with 10 variants compared to the ASFV strain first introduced into Germany in September 2020.

Epidemiological simulation of the spread and inheritance of virus variants illustrates the clustered occurrence of stochastic, geographically distinct variants in a wild boar population without any selection forces. The newly identified characteristic mutation sites were used as genetic markers to enable genomic epidemiology for the different ASFV outbreak strains in Germany. This allowed us to show the geographical distribution and to track the spread of the different ASFV variants in Germany. Using this technique, we were furthermore able to directly connect the ASFV strains responsible for three outbreaks in domestic pigs to the strains circulating in the wild boar population in the same area. Therefore, for the first time since the spread of ASFV GTII in Europe and Asia the transmission pathway between wild and domestic suids was unravelled and spread of ASFV variants can be differentiated in space and time. However, it also highlights the fact that the continuous generation of ASFV WGS is essential, and the only basis on which molecular epidemiology with genetic markers can be performed.

ASFV whole-genome sequencing is laborious and technically challenging, but we were able to generate 22 German and 5 Ukrainian ASFV whole-genome sequences using Illumina based sequencing techniques allowing for single-base resolution and single nucleotide variant identification. The Illumina technology is well suited for ASFV whole-genome sequencing, but the correct calling of G/C homopolymer regions and sequencing the inverted terminal repeats is still error-prone and these regions are therefore excluded from the analyses. To validate the results and rule out sequencing or bioinformatic artefacts, all identified mutation sites in German ASFV sequences have in addition been validated by PCR amplification and Sanger sequencing confirming the whole-genome sequencing results. Therefore, all analyses concerning variant detection and genomic epidemiology are based on validated and confirmed sequencing data.

Interestingly, variant III and IV show genetic variations within four MGF genes e.g. MGF360-10L, MGF360-15R, MGF100-3L and MGF505-4R, while variant II only shows a variation in ASFVs thymidylate-kinase (A240L), an enzyme involved in nucleotide metabolism [14]. Although no function is known for any of the affected MGF genes and corresponding proteins, other ASFV MGF360 and MGF505 genes were shown to be involved in virulence and pathogenicity for example interfering with the hosts interferon response [14,37–39].

The main question remains why this huge increase in the detection of ASFV variants was demonstrated for the first time with German sequencing data. It can be argued that the world-wide number of sequenced samples, especially due to the high efforts needed to generate high-quality ASFV WGS, was not sufficient to cover the extent of ASFV GTII genomic diversity circulating in suids over the past decades. However, the comparison of the German WGS with five Ukrainian and 20 publicly available high-quality ASFV WGS from all over the world draws a different picture. Despite a general tendency to accumulate point mutations over time seen in all ASFV GTII sequences (either silent or leading to single amino-acid switches), a dramatic increase in the detection of High-Impact mutations leading to a relevant genetic frameshift or truncation can be observed in the German ASFV sequences. The data does not comply with the hypothesis of equal mutation dynamics in German virus population and strains previously observed.

The comparison of different variant-sample ratios from different virus populations does not give reliable support to assume that the dynamics of variant generation is constant across affected wild boar populations in Europe. Accelerated variant generation dynamics was suggested when comparing very early (Baltics) and recent (Germany) genomic survey data. Given the assumption that there is an inevitable link between number of emerging variants and the generally increased mutation activity, then the WGS data from Germany in comparison to data from other regions in Eastern Europe indeed suggest an increased mutation rate in the ASFV affected region in Germany and the directly connected region of Western Poland.

However, the increased identification of high-impact mutations in the Polish-German border region may be due to certain selection pressure (Figure 2); or in the other regions the rate of variant emergence is just underestimated due to capacity limits in producing ASFV WGS. To address these uncertainties more high-quality ASFV WGS are needed, especially from Western Poland, where extreme high numbers of ASFV cases have been reported.

The increased mutation rates among German ASFV variants can likely be linked to the HI-mutation in the ASFV DNA PolX gene (O174L), which is shared by all ASFV WGS from Germany as well as the available sequences from western Poland [20,21]. As reported in previous studies, ASFV PolX is a repair polymerase that participates in viral base excision repair, to exchange single damaged nucleotides [33–35]. It therefore seems reasonable to hypothesize that the frameshift mutations in the C-terminus of PolX has a negative effect on its repair activity, thus leading to increased accumulation of mutations in the viral genome. However, despite its function as a repair polymerase, even the wild-type enzyme introduces an unusually high number of errors in its DNA substrates, which has already in the past led to speculations that wild-type PolX might be a strategic mutagenase [40]. This raises the question whether the increased mutation rate is indeed caused by a reduction or perhaps even a gain of activity in the mutated PolX enzyme. While the exact fidelity - e.g. the frequency with which wild-type PolX introduces wrong nucleotides - is still under debate, it is clear that errors are strongly biased towards dG:dGTP misincorporation [40,41]. If such dG:dGTP misincorporation was the reason for the accelerated evolution in German ASFV variants, we would expect to observe a high frequency of dG -> dC and dC -> dG mutations. Yet, no such mutations are found in our dataset (Table 1). This observation goes in line with the previous finding that experimental mutation of the 5’-phosphate binding pocket of PolX, which is also impacted by the frameshift mutation in the German variants, has an even stronger negative effect on dG:dGTP misincorporation efficiency than on Watson-Crick-paired incorporation [34]. It is therefore plausible that the higher mutation rate in German ASFV variants is, at least in part, the result of overall reduced enzymatic activity rather than increased dG:dGTP misincorporation efficiency of the reparative polymerase PolX.

## Conclusion

In conclusion, we report here the emergence of distinct ASFV variants that point to a higher sequence variability of ASFV in strains observed at the German-Polish border. We identified a frameshift mutation in the O174L gene/ PolX that affects the 5’binding pocket of the enzyme as plausible cause. The resulting ASFV variants allow, on the upside, for the first time a meaningful genomic ASFV epidemiology. On the downside, the accelerated occurrence of viral variants has the potential to result in ASFV variants with novel features which might in the future dramatically influence the course of the ASFV epizootic with unknown outcome.

## Acknowledgements

We gratefully thank Patrick Zitzow, Ulrike Kleinert and Robin Brandt for excellent technical assistance. Olha Chechet and Mykola Sushko for the selection of representative samples and providing sample material.

